# Sieve tube structural variation in *Austrobaileya scandens* and its significance for lianescence

**DOI:** 10.1101/2021.09.23.461614

**Authors:** Juan M. Losada, Zhe He, N. Michele Holbrook

## Abstract

Lianas are characterized by large leaf areas and slender stems, a combination of features that require an efficient vascular system. The only extant member of the Austrobaileyaceae, is an endemic twining liana of the tropical Australian forests with well-known xylem hydraulic traits. However, the vascular phloem continuum through aerial organs remains understudied.

We analyzed the structure of phloem conduits across leaf veins and stems of *A. scandens*, combining topological data obtained through light and electron microscopy, with current models of phloem transport.

Leaves displayed a low xylem to phloem ratio compared with leaves of other angiosperms, with vascular elements invariant in diameter along the midrib, but tapered across vein hierarchies. Sieve plate pore radii were extremely small: 0.08 µm in minor veins, increasing to 0.12µm in the petiole and only to 0.20µm at the base of the stem, tens of meters away. Searcher branches contained tube shaped phloem conduits with a pectin-rich wall, whereas twining stems displayed sieve elements with tangential connections that displayed a greater fraction of the tubes populated with an astonishing number of sieve plates.

Hydraulic segmentation of the leaves in Austrobaileyaceae correlate with vesseless leaves that benefit photoassimilate export through volumetric scaling of the sieve tube elements. Yet, compared with canopy dominant trees, the geometrical properties of the sieve tube in twining stems, restrict considerably energy distribution in the sub-canopy layers, potentially favoring the allocation of assimilates toward the elongating branches. Thus, the conductive xylem of twining stems contrasts with a poorly conductive phloem that meets the mechanical constraints of lianescence.

## INTRODUCTION

Lianas constitute about 40% of the woody individuals in tropical forests and make an outsize contribution to forest productivity and response to disturbance (Gerwing and Farias, 2000; Schnitzer et al., 2012). Lianas elevate a dense leaf canopy tens of meters aboveground with a minimal investment in self-support (Darwin, 1868; Baillaud, 1962). Anatomical features that allow lianas to meet the evaporative demands of their leaves despite having slender stems include large diameter xylem vessels, and thus greater hydraulic efficiency compared with that of co-occurring trees and shrubs (Carlquist, 1991; Ewers et al., 1991; Isnard et al., 2003; Isnard and Silk, 2009; Wyka et al., 2013; Chen et al., 2014; Pace et al., 2015, 2018; Chery et al., 2020). The allometrical demands of the climbing habit might be expected to extend to the phloem. Yet, the long-distance effectiveness of carbohydrate transport remains poorly documented in lianas, due to the scarce information on the intersection between the vascular structure of the phloem and lianescence.

Previous anatomical descriptions of the stems of large bodied vines belonging to Loganiaceae, Malpighiaceae, Bignoniaceae, Fabaceae or Sapindaceae emphasize novel topologies with distinctive cambia: anomalous phloem architectures that evolved with the scandent habit and represent traits that associate with both higher flexibility and resilience to damage (Ewers and Fisher, 1991; Fisher and Blanco, 2014; Moya et al., 2017; Pace et al., 2011; 2015; 2018; Chery et al., 2020). Only a handful of works address structural variation of the sieve tube elements in relation to axial transport in lianas. Other woody life forms such as trees and shrubs are better characterized, and scale the geometry of phloem conduits, pore sizes, and sieve plate number with stem diameter (i.e. distance) (Liesche et al., 2017; Savage et al., 2017; Losada and Holbrook, 2019; Clerx et al., 2020; Barceló-Anguiano et al., 2021a,b). In turn, sieve tube geometry varied little axially in the long and slender stems of *Ipomoea* vines, but sieve pore sizes connecting conduits enlarged from the top to the base of the stem, thus allowing transport at long distances, in agreement with the pressure-flow hypothesis developed by Münch in 1930 (Knoblauch et al., 2016).

*Austrobaileya scandens* is a large-bodied and long-lived liana native to Queensland (Australia), which reaches more than 15m aboveground by twining around trees as its only support (Bailey and Swamy 1949). *A. scandens* exhibits low photosynthetic rates and a slow stomatal response to drops in environmental humidity (Feild et al., 2003a,b; Feild and Arens, 2005, 2007; Feild and Wilson, 2012; Barral et al., 2013). Anatomically, the stems of *A. scandens* display wide xylem vessels (Carlquist, 2001) and a secondary phloem with extremely angled compound sieve plate (Behnke, 1986). This sieve tube morphology, which resembles the phloem of gymnosperms, drove early speculations on a possible transitional tissue toward the `pipe-like’ sieve tubes of most angiosperms (Bailey and Swamy, 1949; Srivastava, 1970; Behnke, 1986), although angled plate could simply compensate for the small plate pores. Alternatively, the peculiar morphology of the secondary phloem of *Austrobaileya* may be related to maintaining the functional demands of the phloem as climbing stems undergo mechanical deformation. To better understand vascular transport in large-bodied lianas, we studied the phloem of leaves and stems, so far missing in this species. The main goal of this study was to understand the structure of the phloem in relation to 1) axial transport of carbohydrates from the sites of photosynthesis through the slender stems; 2) mechanical demands of climbing.

## MATERIALS AND METHODS

### Plant material

We used the germplasm resources of the Arnold Arboretum of Harvard University, which includes greenhouse-grown *Austrobaileya scandens* plants ranging in age from 20 to ∼ 25 years old. The leaf area of seven terminal branches from different plants was measured by photographing each leaf from the branch tip to the seventh node (total n=84 leaves (or nodes?)). Since opposite leaves displayed similar areas, their average at each node was used as a measurement of leaf area up to the seventh node, where leaf area reached a maximum.

### Leaf anatomy

Five mature leaves were used to determine the area occupied by the vascular elements of the leaf midrib: consecutive hand transverse sections were cut with a micro scalpel at three different positions in the leaves: the petiole, the mid part of mid vein, and about 2cm back from midrib tip. Each section was stained with 0.1% w/v aniline blue in PO4 K3 (pH 10) (Linskens and Esser, 1957), observed with a Zeiss Axiophot epifluorescent microscope using the DAPI narrow filter band (excitation 365 nm, bandpass 12 nm; dichroic mirror FT 395 nm; barrier filter LP397 nm), and photographed with an AxioCam 512 Color linked to the AxioVision software (Zeiss, Oberkoche, Germany). The cross-sectional area of xylem and phloem was measured (totaling 45 measures for each tissue) and, for the 27 cross sections in which all vascular elements were visible, the number of vascular elements in each tissue were counted

Five additional leaves were selected for measurements of individual sieve tubes and xylem vessels. Longitudinal sections of the petiole, midrib, second, third, and fourth order veins were obtained with a scalpel, placed in a solution containing acetic acid: hydrogen peroxide 1:1 v/v, left incubating at 60ºC for two days, and then mounted onto glass slides for microscopy observation and images obtained (n>250 total vessel elements measured). Similar areas were used to obtain fresh longitudinal sections (less than 1mm thick), mounted onto slides, stained with 0.1% aniline blue, counterstained with calcofluor white for cellulose (Hughes and McCully, 1975), photographed and measured (at least 35 sieve tubes per vein order, n=250).

### Stem anatomy

Branches were classified according to both their diameter and stiffness (see Fig. 1): < 2 mm (primary growth, flexible), < 2 mm (stiff and twisted around a support), 6-10 mm (green shoots with secondary growth), 11-15 mm (stems at the base of the vine with highly twisted-irregular bark). First, we evaluated the general anatomy of the stems in transverse sections at the top three axial locations (the widest stems could not be cut transverselly to preserve the plants): flexible searchers (2mm diameter), stiff thin branches (2mm diameter), and mature stems (6-10mm diameter). Stem material was maintained in 1X Tris-buffered saline (TBS), sectioned transversally at 50 μm with a Reichert-Jung Hn-40 sliding microtome (Austria), mounted onto glass slides, and stained with aniline blue that stains callose of the sieve tubes, or with 0.1% of acridine orange in TBS buffer, which yields fluorescence of the lignified tissues (Robertson et al., 1992). To sample the phloem tissue, the external side of stems containing the bark were extracted with a sharp knife at the different positions described above, kept in buffer, sectioned longitudinally with a microscalpel, and stained with aniline blue. We measured the length and width of at least 30 sieve tubes per sampling location (total n=107), and the number of sieve areas per compound plate (total n=45).

**Figure 1.**
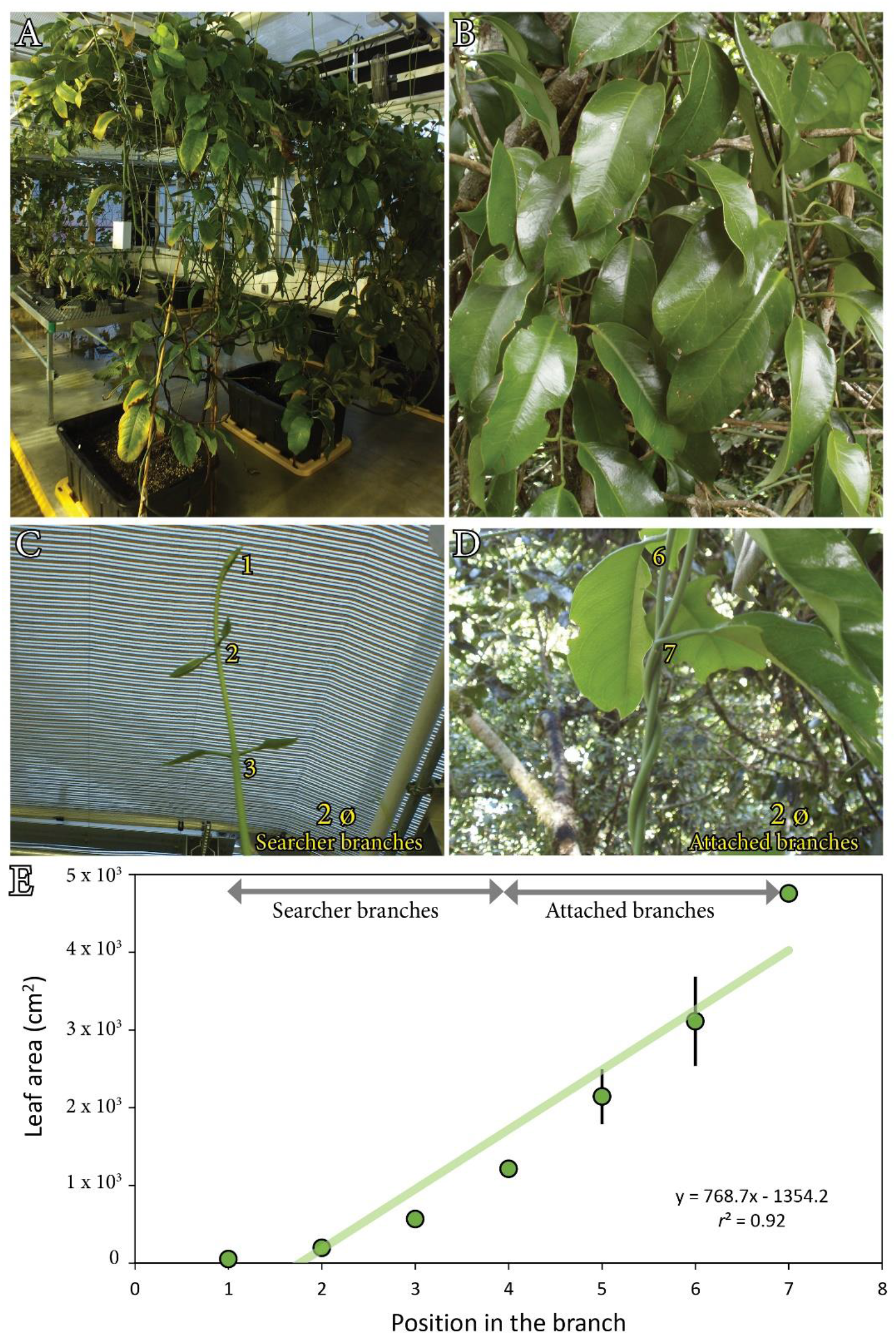
**A**. Adult vines of *Austrobaileya scandens* in the greenhouse. **B**. *A. scandens* vines at the canopy layer in the tropical rainforest of Queensland (Australia). **C**. Apical part of a ‘searcher’ branch in the greenhouse. **D**. Branches twisted around a stable support without increasing their diameter. **E**. Leaf area at each node increases basipetally following a linear fashion at a *p*<0.05.

### Scanning electron microscopy

To evaluate the size of pores that make up the sieve plates, samples of all leaf veins and stem axial areas (see above description) were cut, frozen in liquid nitrogen, and transferred to super-chilled ethanol, prior to being sectioned in different angles with respect to the main axis with the goal of obtaining sections aligned with the highly angled sieve plates. These sections were incubated within a mixture of 0.1% w/v proteinase K dissolved in 50 mM Tris-HCl buffer, 1.5 mM Ca^2+^ acetate and 8% Triton X-100, pH 8.0 (Mullendore et al., 2010), in a water bath at 60°C for 2 weeks. After rinsing with ethanol once, and with distilled water thrice, the sections were incubated with a 1% w/v aqueous solution of α-amylase for 2 days at 60°C, then rinsed thrice in water and finally freeze-dried for 24 h with a Freeze Dyer (Labconco Freeze Dry System). Samples were then mounted on SEM studs and sputter coated with gold-palladium using a Denton Vacuum Desk II Sputter Coater for 180 s at 20 V and 6.67 Pa. Samples were imaged with a JEOL-6010LV scanning electron microscopy (SEM) (JEOL, Peabody, MA, USA), using high vacuum and an accelerating voltage of 10–15 kV. The size of sieve pores, of sieve areas, and pore density were measured. Although sieve pore radius was measured for each vein order, due to the small size of the sieve plates within higher order veins, the preservation of intact sieve plates was poor. As a result, we were only able to determine sieve plate size and pore density for the midrib (n=5). The total number of pores measured was large (n=105 for petiole; n=262 for primary vein; n= 84 for minor vein orders). In the thinnest stems, the stiffness of the external fiber layer, along with the fragility of the sieve tubes, limited preservation –and thus visualization-of the pores, relative to the base of the stem where preservation was much better and resulted in n=600 pores measured.

### Phloem sap velocity in *Austrobaileya scandens*

Phloem transport velocity was measured by tracking the movement of the fluorescent dye esculin (reviewed by Knoblauch et al., 2015) in the secondary veins of a five-year-old vine at 11:00 h for several days. A young plant (five years approximately) with fully formed leaves, was watered to saturation, and then one of the long branches immobilized on the microscope stage, with the abaxial surface exposed upward. A 50 μL droplet of 5mM aqueous esculin mixture, combined with 0.1% of the SilEnergy surfactant (RedRiver Specialties, Shreveport, LA, USA), was applied with a micropipette to the junction of the second with minor order veins (adapted from Jensen et al., 2011, Savage et al., 2013). To allow better permeabilization of the dye, the tip of the pipette was used to slightly abrade the thick cuticle, and a small window was opened, and was covered with the liquid during the experiment to prevent desiccation. To track the movement of the dye, we used a portable Stereo Microscope Fluorescence Adapter with 510–540 nm excitation wavelength, and a long pass 405 UV nm filter band (NIGHTSEA, Lexington, MA, USA). Time-lapse images were obtained every 10 s for 30min with a Zeiss v12 dissecting microscope using the 0.63× PlanApo objective and an AxioCam 512 Color camera connected to the AxioVision software. Despite multiple attempts, the movement of the dye could only be observed in two leaves. In both cases, fluorescence increased gradually downstream the vein a few minutes after the dye was applied. Velocity was measured by thresholding the images (Hue 119-188; Saturation: 97-255; Brightness: 0-255nm), cleaning the dark outliers with less than 2 pixels’ radius, and then calculating the time elapsed by pixel accumulation two centimeters away from the area where the dye was applied (n=2).

### Immunolocalization of a branched pectin epitope

Immunolocalization of branched 1-4 galactan with the LM26 monoclonal antibody (PlantProbes, Leeds, UK), was performed in the searcher stems (stiffer stems were poorly embedded and difficult to section). Segments of 0.5cm from flexible branches were fixed with 4% w/v acrolein (Polysciences, Warrington, PA, USA) in a modified piperazine-N, N′-bis (2-ethanesulfonic acid) (PIPES) buffer adjusted to pH 6.8 (50 mM PIPES and 1 mM MgSO4 from BDH, London, UK; and 5 mM EGTA) for 24 h, then washed and dehydrated through a series of increasing aqueous acetone concentrations, one hour each: 10%, 30%, 50%, 70%, 90%, 100%. After that, they were incubated in a solution containing the Technovit 8100 for two weeks, and then hardened in anoxic conditions at 4ºC. Longitudinal and transverse 4µm sections were obtained with a Leica EM UC7 ultramicrotome (Leica Microsystems, Wetzlar, GE), mounted onto superfrost slides, and then used for immunolocalization. Pre-incubation with 5% bovine albumin serum (BSA), and three washes in 1XPBS, were followed by incubation with the LM26 monoclonal antibody for 1h. After washing the primary antibody with PBS, an anti-rat alexa488 secondary antibody, with a fluorescein isothianate marker-FITC was applied for 1h. Samples were then washed and observed with a Leica DM2500 microscope equipped with epifluorescence and a Leica DM600 camera, combining the 405 filter for autofluorescence with the 488 filter for the specific signal of the FITC.

### Image analysis and statistics

The transverse areas of the leaf vascular tissues were manually outlined, individual tube number counts and length/width measurements evaluated on images with the Image J 1.51d software (National Institutes of Health, Bethesda, MD, USA). Individual images from stem cross sections were aligned and merged into a composite image with the Photoshop software (Adobe Systems, Newton, MA, USA).

Averages from geometrical evaluations were compared using a one-way ANOVA and the post hoc Tukey test at a *p*<0.05, using the SPSS software for statistics.

## RESULTS

### General anatomy of *Austrobaileya scandens*

*A. scandens* are evergreen vines with long stems with opposite leaves and branches (Fig. 1A,B) see also Bailey and Swamy 1949; Metcalfe 1987; Feild et al., 2003b). The self-supported young apical branches were cylindrical, flexible, and about 2mm diameter (Fig. 1C). These young stems hardened around 10cm back from the tip, acquiring an elliptical shape, coincident with their coiling around a solid support (Fig. 1D). Despite the minimal variation of branch diameter, a linear gradient of leaf expansion was noticed basipetally, with younger leaves at the flexible branch tips, and mature ones farther away in the coiled branches (Fig. 1E).

### Vascular geometry in the leaves of *Austrobaileya scandens*

Mature leaves of *A. scandens* (Fig. 2A) were coriaceous, with a short petiole and an entire lamina. The vasculature of the petiole included adaxial xylem tissue composed mainly of squared tracheids, and abaxial phloem composed of sieve tubes (identified by the presence of callose), rays, and parenchymatous tissue (Fig. 2B). While fiber caps were lacking in the petiole, a massive protective perivascular fiber cap proliferated in the midrib (Fig. 2C-E). The cross-sectional areas of both the xylem and the phloem were greatest in the petiole and narrowed linearly toward the tip of the major vein [xylem: y=-0.05x+0.20(mm^2^), phloem: y=-0.04x+0.15(mm^2^)]. Strikingly, conduit diameter of both phloem and xylem were invariant along the midrib, but the number of conduits decreased by a factor of 2.6 from the petiole to the midrib tip.

**Figure 2.**
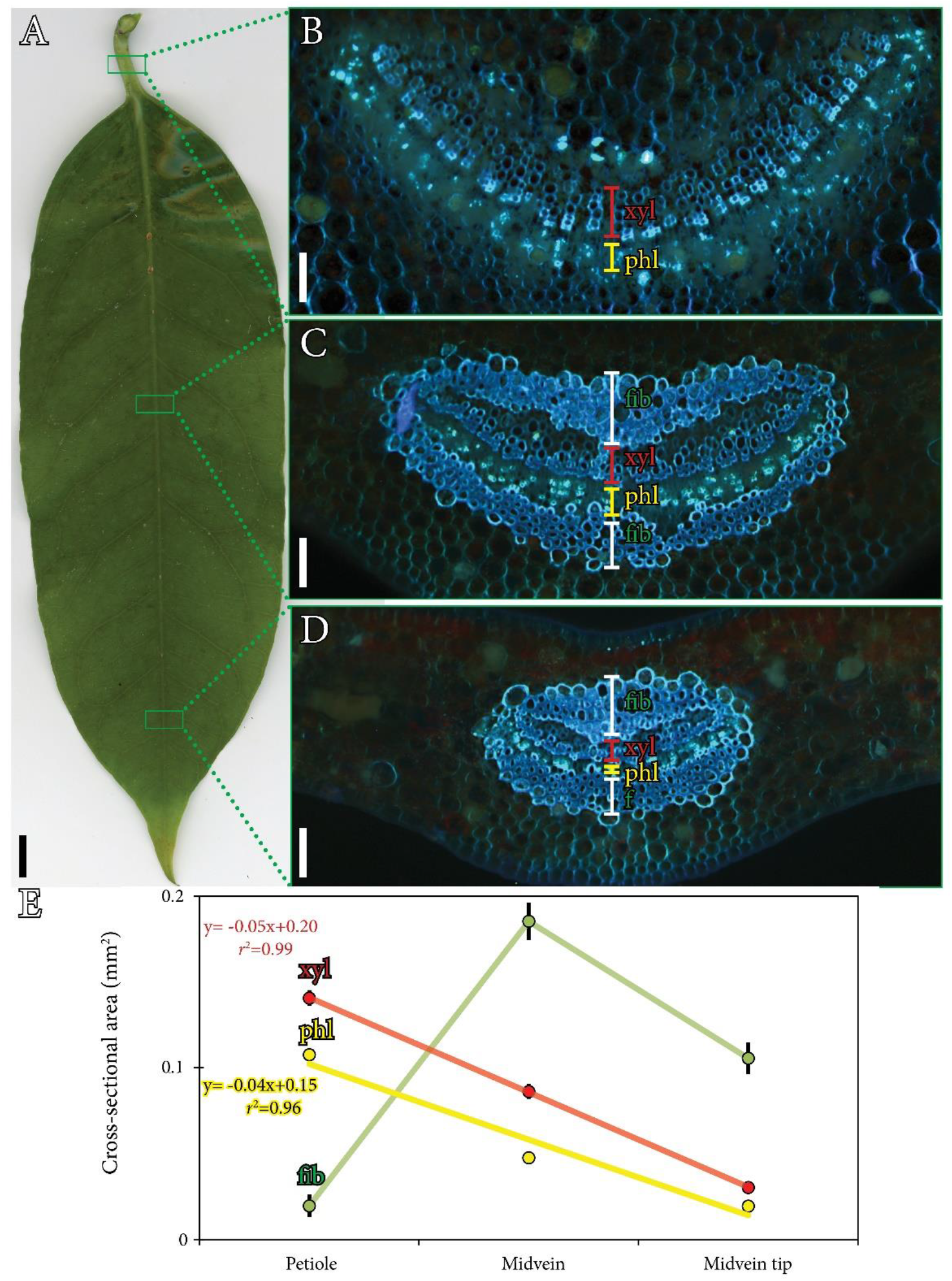
Vascular anatomy of the continuum petiole-mid vein of *Austrobaileya scandens* leaves. **A**. Mature leaf of A. scandens. **B**. Vascular tissues of the fiber less petiole. **C**. Vasculature in the mid part of the major vein showing a massive layer of pericyclic fibers surrounding the xylem and the phloem. **D**. Vasculature at the tip of the mid vein. **E**. Cross sectional areas of the vascular tissues at the three positions in the midvein: while the cross-sectional areas of xylem (red) and phloem (yellow) decreased linearly toward the tip (*p*<0.05), the fibers (green) occupied a wide cross-sectional area in the leaf lamina, but not in the petiole. **B-D**. Cross sections stained with aniline blue to detect callose of the sieve tube elements. fib, fibers; phl, phloem; xyl, xylem. Scale bars A: 1cm; B-D: 200 µm.

Xylem and phloem conduits varied in size across vein hierarchies (Fig. 3A), gradually decreasing in size from major to minor veins, except in the petiole, where tracheids were shorter and narrower than those of the midrib (Fig. 3B). Although sieve tube elements were also shorter in the petiole, they slightly increased in diameter with respect to midrib (Fig. 3C). Pore sizes of the sieve elements decreased their radii from 0.12µm in the petiole to less than 0.08µm in the minor veins (Supplementary figure 1). Functionally, foliar phloem architecture correlated with a low bulk velocity of the mobile dye tracer esculin hydrate *in vivo* (Fig. 4; Video S1), revealing an average rate of 11µm s^-1^ in the second order veins (n=2).

**Figure 3.**
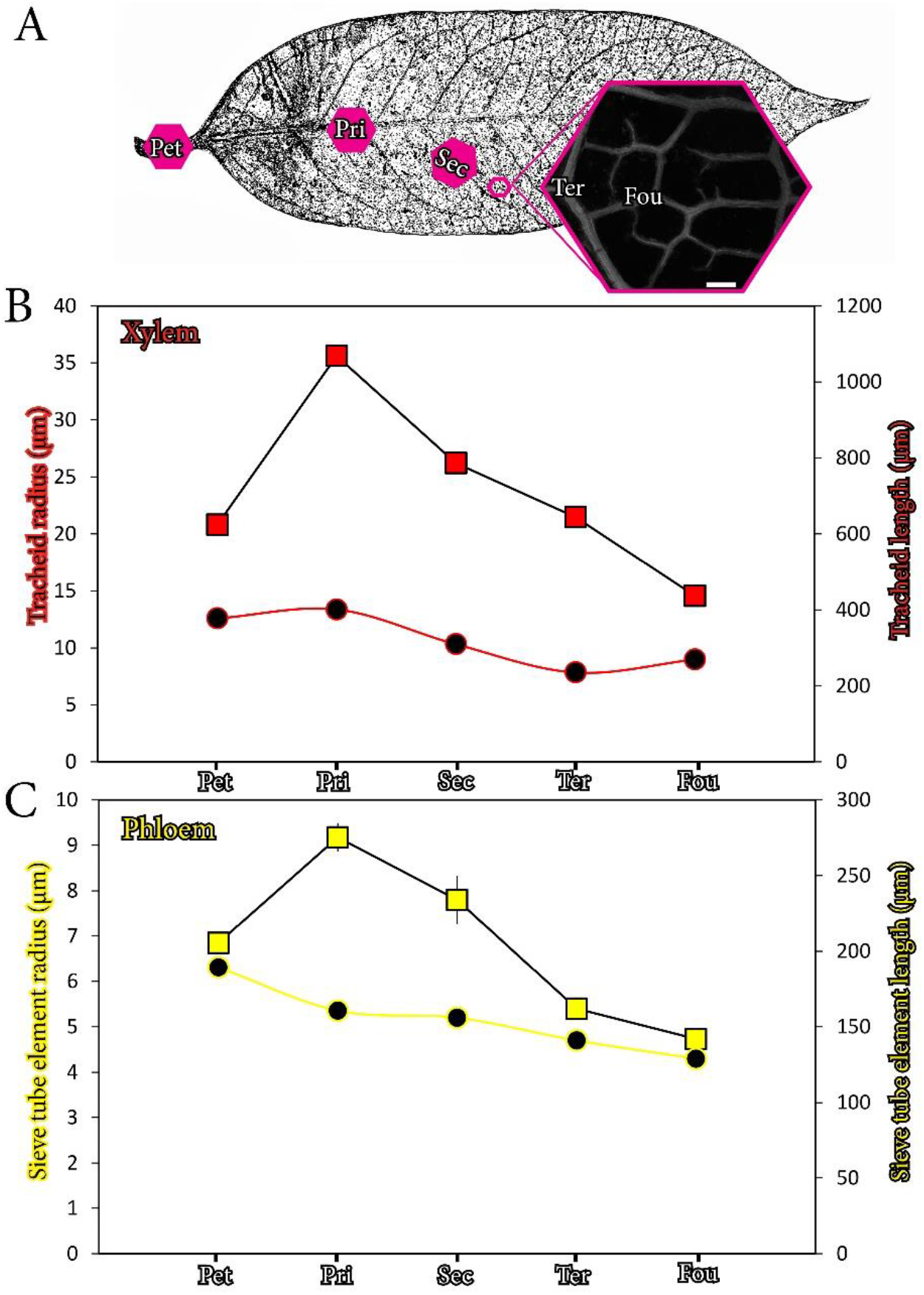
Geometrical scaling of the vascular elements in the leaves of *Austrobaileya scandens*. **A**. General view of the leaf veins showing the sampling areas (pink hexagons) and closeup of the minor veins. **B-C**. Length (squares) and radius (circles) of the individual vessel elements (red), and sieve tube elements (yellow) across vein orders; error bars represent standard error at a *p*<0.05. Inset in panel **A** shows a fluorescence image section of the leaf stained with the Feulgen reactive, cleared, and cuticle dissected. Fou, fourth order veins; pet, petiole; pri, primary vein; sec, secondary vein; ter, tertiary vein. Scale bar: 500µm.

**Figure 4.**
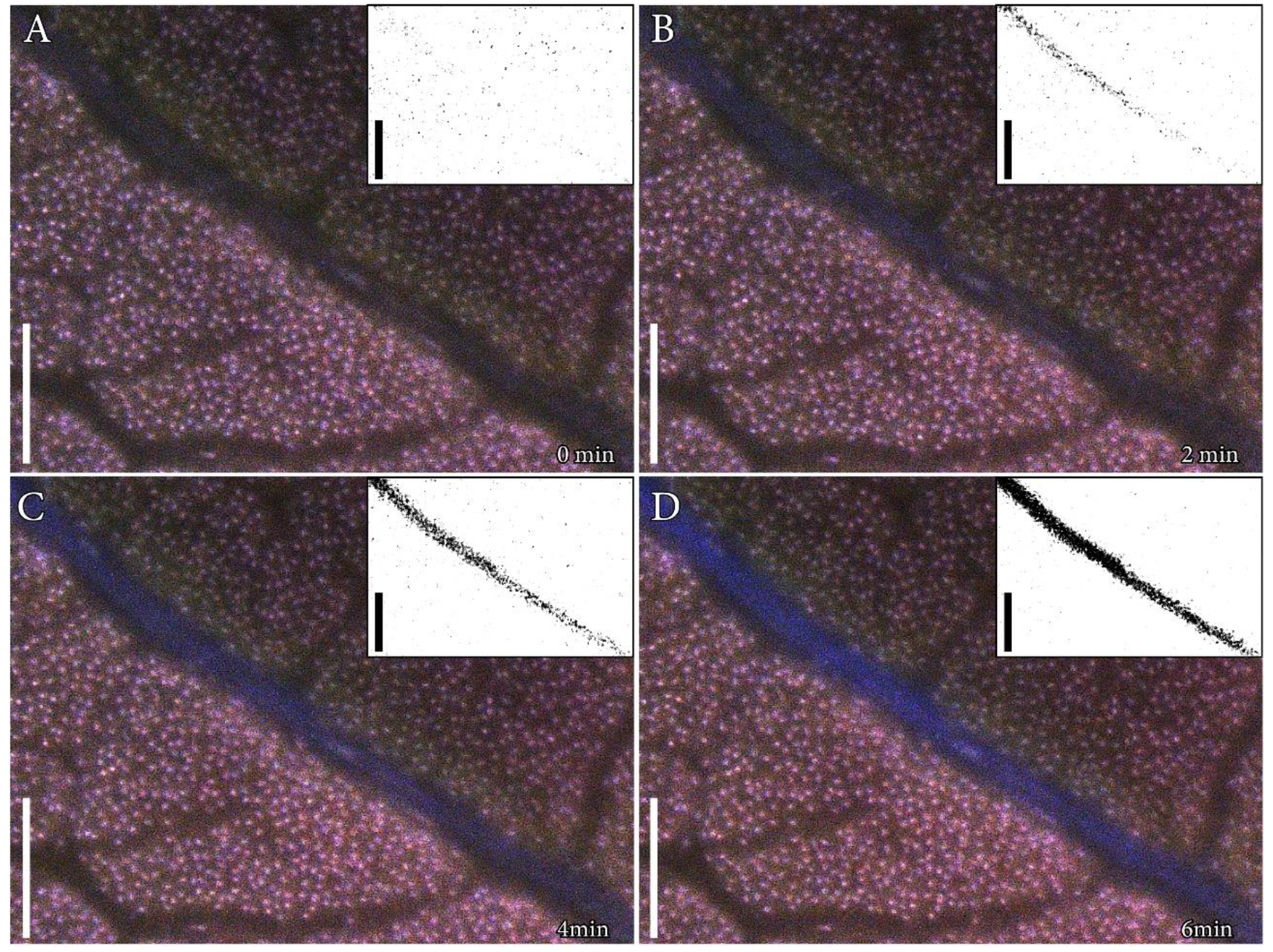
Velocity of the phloem sap in *Austrobaileya scandens* leaves. Time-lapse images of a secondary vein showing a 2 min advancement of esculin hydrate dye from 0min (A) to 6min (D). Insets show the thresholding applied to the sequential images for the calculation of velocity, with a gradual accumulation of the dye within the veins. Scale bars: 1000μm.

### Ontogeny of the phloem in the stems of *Austrobaileya scandens*

Cross sections of stems at different axial positions revealed ontogenic differences in the eustele, which correlated with stem mechanical properties. In the flexible searcher stems, a large pith was surrounded by discontinuous primary xylem traces consisting of a few tracheids (Fig. 5A), but a more continuous phloem layer (Fig. 5B). Twining correlated with a sharp increase in the stiffness due to lignification of the pith and the development of an extra phloem pericyclic fiber layer (Fig. 5C). In wider stems (the ones that traverse longest distances), the central pith constituted the largest fraction of the cross-sectional area, and the axial secondary phloem displayed a lobed morphology (Fig. 5D-E), with sieve tubes closer to the vascular cambium and separated from the pericyclic fibers by parenchyma (Fig 5F).

**Figure 5.**
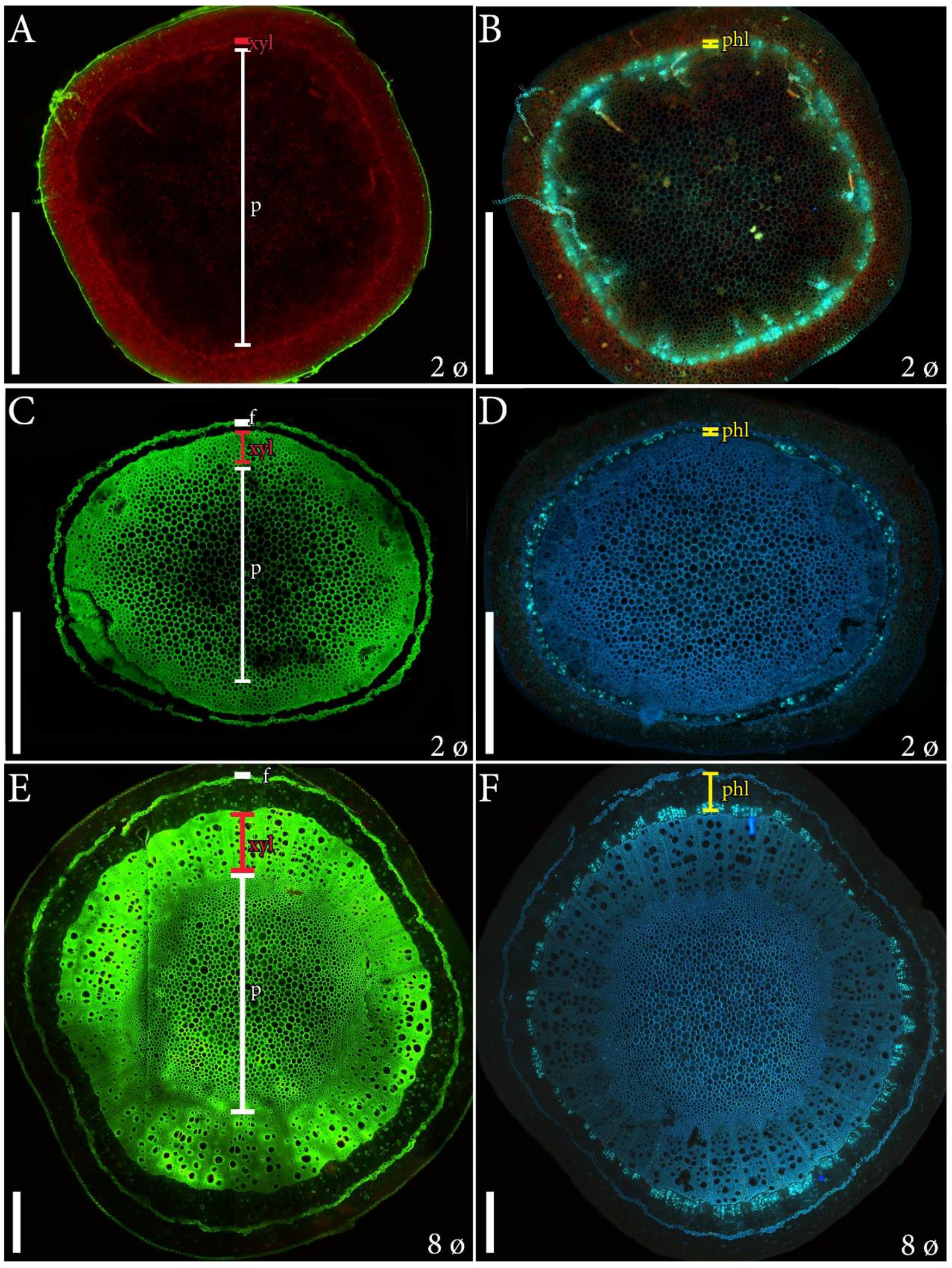
Anatomy of the *Austrobaileya scandens* stems. **A**. Cross section of a 2mm diameter ‘searcher’ branch tip with primary growth, with an extensive central pith. **B**. Same stem showing callose in the continuous phloem ring (fluorescent green). **C**. Cross section of a stiff 2mm diameter branch showing a dramatic increase in lignification of the pith, the xylem, and a pericyclic fiber cap (fluorescent green). **D**. The phloem forms a thin layer between the xylem and the fiber cap. **E**. Wider stems (8mm diameter) with a high degree of lignification in all central tissues, wide vessels of the xylem, and enlarged areas between the xylem and the fiber cap. **F**. The fascicular phloem tissue was separated by axial multilayered phloem parenchyma and the active tubes were in the vicinity of the vascular cambium. **A**,**C**,**E**, cross sections of stems stained with acridine orange, which displays fluorescence of the lignified tissues; **B**,**D**,**F**, cross sections of the stems stained with aniline blue for callose in the sieve tube elements of the phloem. F, fibers; p, pith; phl, phloem; xyl, xylem. Scale bars: 1mm.

In line with the demands for climbing and the need for flexible searcher branches, a galacturonan-rich pectic epitope localized in the sieve tube walls of the searcher stems of *A. scandens* (Fig. 6). While numerous cells composed the stems in cross section (Fig. 6A), this epitope specifically showed the scattered distribution of sieve tubes (Fig. 6B, C). Upon closer inspection, the signal of the antibody corresponded with the areas of the sieve tubes between the plasma membrane and the cell walls (Fig. 6D-F). In longitudinal section, the sieve tubes were not distinguishable from other phloem cell types (Fig. 6G), but the presence of this epitope highlighted their longitudinal silhouette (Fig. 6H-I), showing slightly tangential sieve plates connections (Fig. 6J-L).

**Figure 6.**
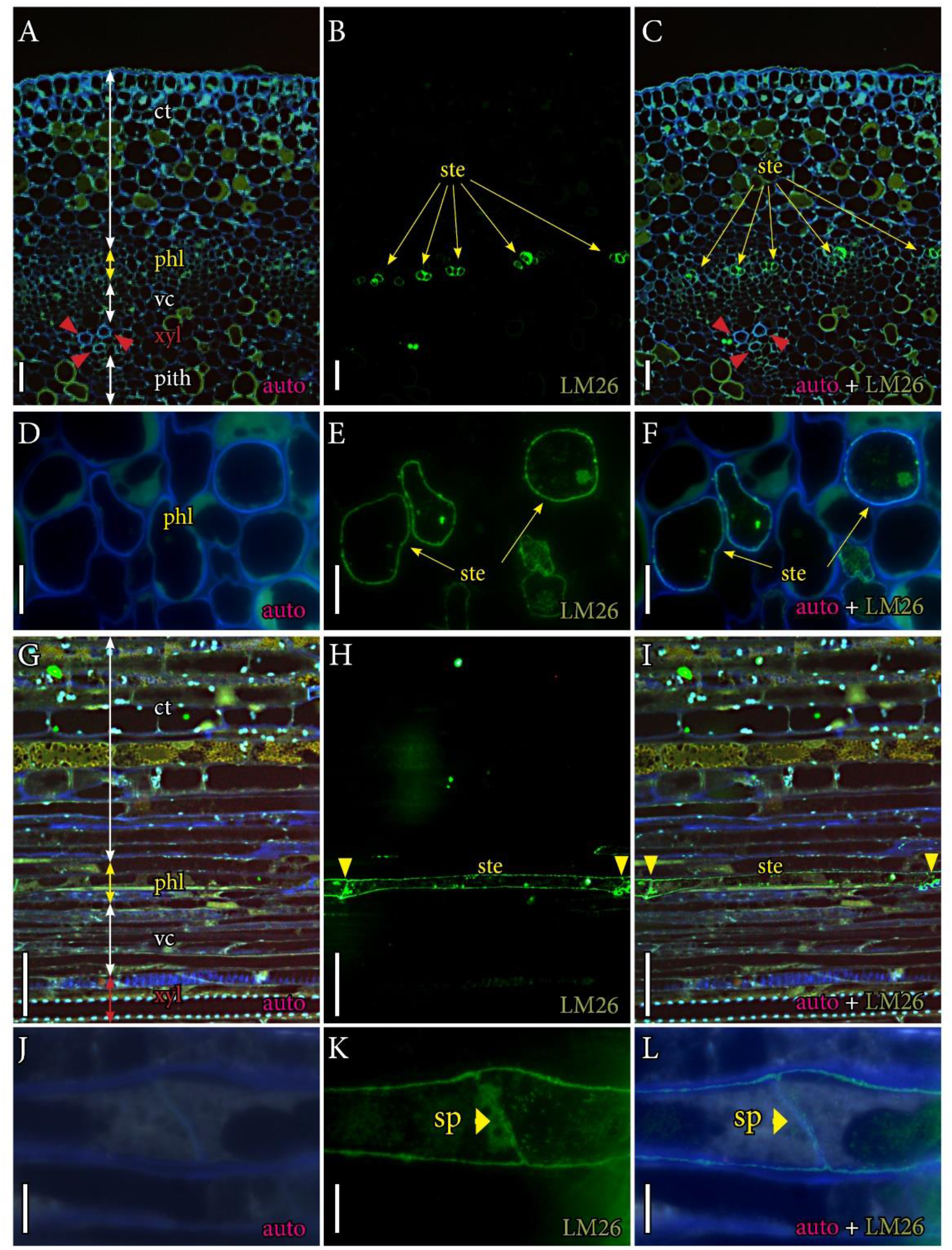
Immunolocalization of LM26 epitope in the sieve tube elements of searcher branches of *Austrobaileya scandens*. **A**. Transverse section showing multicellular arrangement in the different tissue layers, including three isolated tracheids that compose the primary xylem (red arrowheads). **B**. Same section displaying the sieve tube elements of the primary phloem forming a continuous ring (yellow arrows). **C**. Merged images. **D** Close up of the phloem in cross section. **E**. The profile of the sieve tubes in cross section after immunolocalization (yellow arrows). **F**. Merged images. **G**. Longitudinal section of the branch displaying he external part in the top and the internal tissues at the bottom. **H**. Longitudinal profile of the sieve tube element wall after immunolocalization, yellow arrowheads define the connection between tubes. **I**. Merged images. **J**. Detail of the sieve tube connections. **K**. Sieve plate in the primary sieve tubes are slightly tangential (yellow arrowhead). **L**. Merged images. 4µm thick transverse (**A-F**) or longitudinal (**G-L**) sections of the flexible branches displaying autofluorescence with the 405nm filter (**A**,**D**,**G**,**J**), immunolocalized with the LM26 monoclonal antibody (**B**,**E**,**H**,**K**), and merged images (**C**,**F**,**I**,**L**). Ct, cortical tissue; phl, phloem; sp, sieve plate; vc, vascular cambium; xyl, xylem. **A-C** scale bars: 50 µm; **G-I** scale bars: 25 µm; **D-F**, J-L scale bars: 10 µm.

The morphology of the sieve elements changed dramatically from flexible stems to stems supported by twinning, gradually increasing the number of sieve plate connections, as well as their angled position at the end of the conduits (Fig. 7A,B). In the elongated slender stems, sieve elements had a vermiform appearance with extremely angled tangential connections that resulted in an astonishing number of sieve areas (Fig. 7A-C). Numerous small pores (0.20µm on average, n=600), whose radius was similar to those of the leaf petioles, populated the sieve plates (Behnke, 1986). In turn, sieve tube element dimensions increased from thinner to thicker stems, but their length reduced at the base of the vine (Fig. 7E).

**Figure 7.**
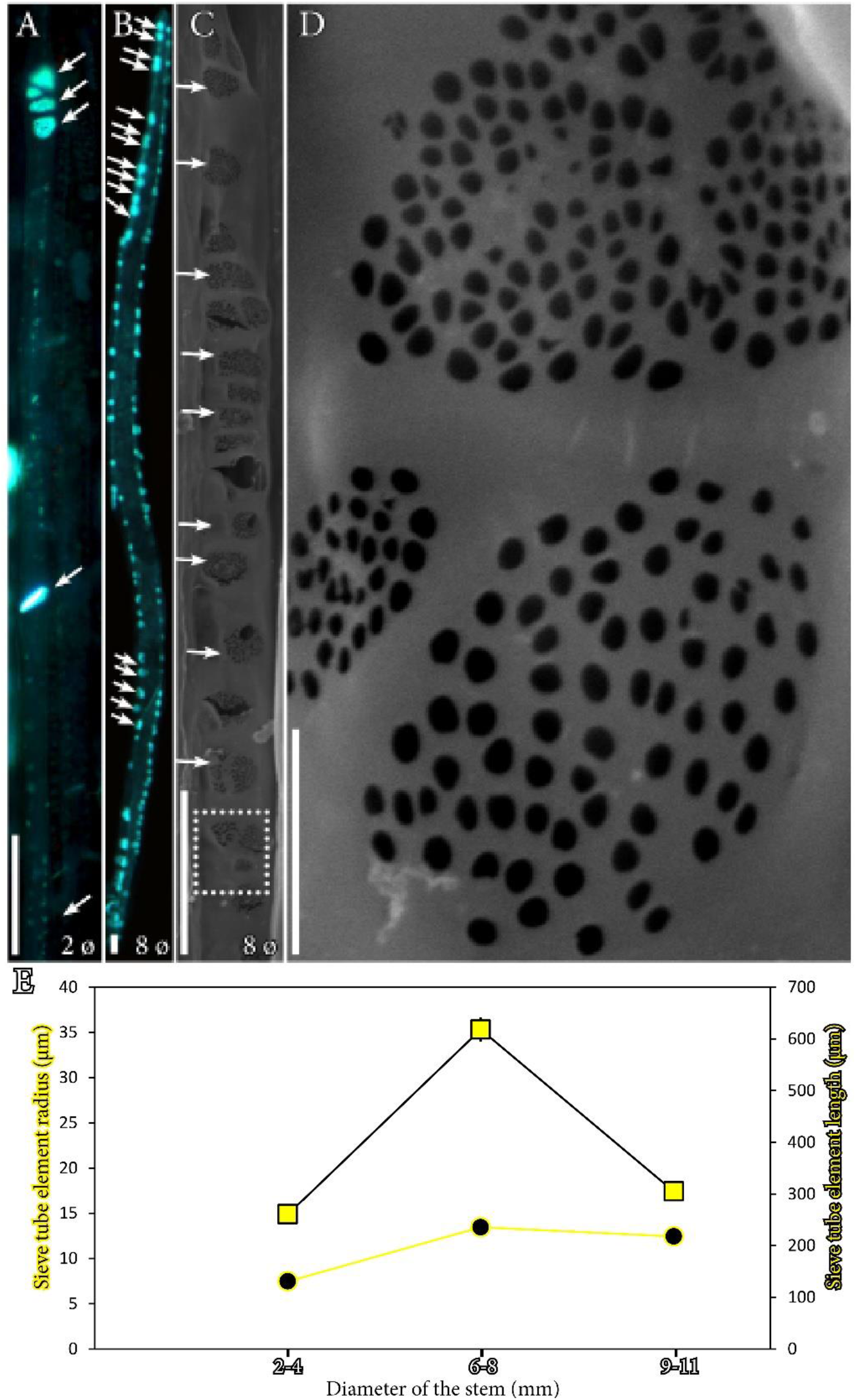
Morphology of the sieve tube elements in the stems of *Austrobaileya scandens*. **A**. Longitudinal view of a sieve tube element from a flexible branch, 2mm diameter, showing either simple or compound sieve plate connections (arrows). **B**. Sieve tube element of an 8mm diameter stem showing numerous sieve plate connections between tubes (arrows). **C**. Longitudinal view of a sieve tube element with scanning electron microscopy displaying numerous sieve plates and sieve areas all along the tube wall (arrows). **D**. Close ups of three sieve plates (analogous to the white dotted square in figure C), with details of pores. **E**. Sieve tube element length (squares), and width (circles) across axial stems of different diameter ranges; bars represent the standard error at a *p*<0.05. **A, B**, longitudinal hand sections of the stems stained with aniline blue to detect callose (bright fluorescence); **C**,**D**, scanning electron microscopy images showing the sieve plates and the sieve pores in detail. A-C scale bars: 20um; D: 2µm.

### Phloem hydraulic resistance from leaves to stems of *Austrobaileya scandens*

The measured velocity (assumed as constant) was combined with anatomical data (average sieve tube radii, and pore radii/length at each location along the transport pathway in both leaves and stems), supposing a continuous pipe-like cylindrical tubing system formed by sieve elements connected in series, as modelled previously for vines and trees (Knoblauch et al., 2016; Savage et al., 2017). The Hagen-Poiseuille equation was the reference to understand the axial conductivity of the sieve tube. Due to the difficulty on measuring the actual length of the twisted stems, we calculated the total resistance per tube length (Pa s m^-4^) at the relative axial positions along the transport path from leaves to the base of the stem. The poor preservation the thinner stems allowed only a few measurements of pore size, which revealed an average radius of 0.19µm, but this was quite similar to the pore radius at the base of the stem (0.21µm), where preservation was much better and resulted in n=600 pores measured. The following assumptions were made for calculations of the hydraulic resistance: (1) sieve areas and pore density were equal to the midrib value for higher vein orders; (2) Pore size at the tip of the stems were similar to those measured at the vine base. The latter assumption is very conservative, and likely underestimates the resistance of this stem areas, given that previous reports showed smaller pore sizes at the tips of vine stems compared with their base (see Knoblauch et al., 2016).

In general, the small pores of sieve plates mainly drove the total sieve tube resistance (Fig. 8). However, while variations of pore size from the petiole to stems were subtle (0.12µm and 0.20µm respectively), both tube volumetric increase and the number of sieve plates per end tube were the main factors allowing a drop in sap flow resistance from leaves to stems. At the base of the stems, sap flow was slightly constrained due to the sieve element shortening.

**Figure 8.**
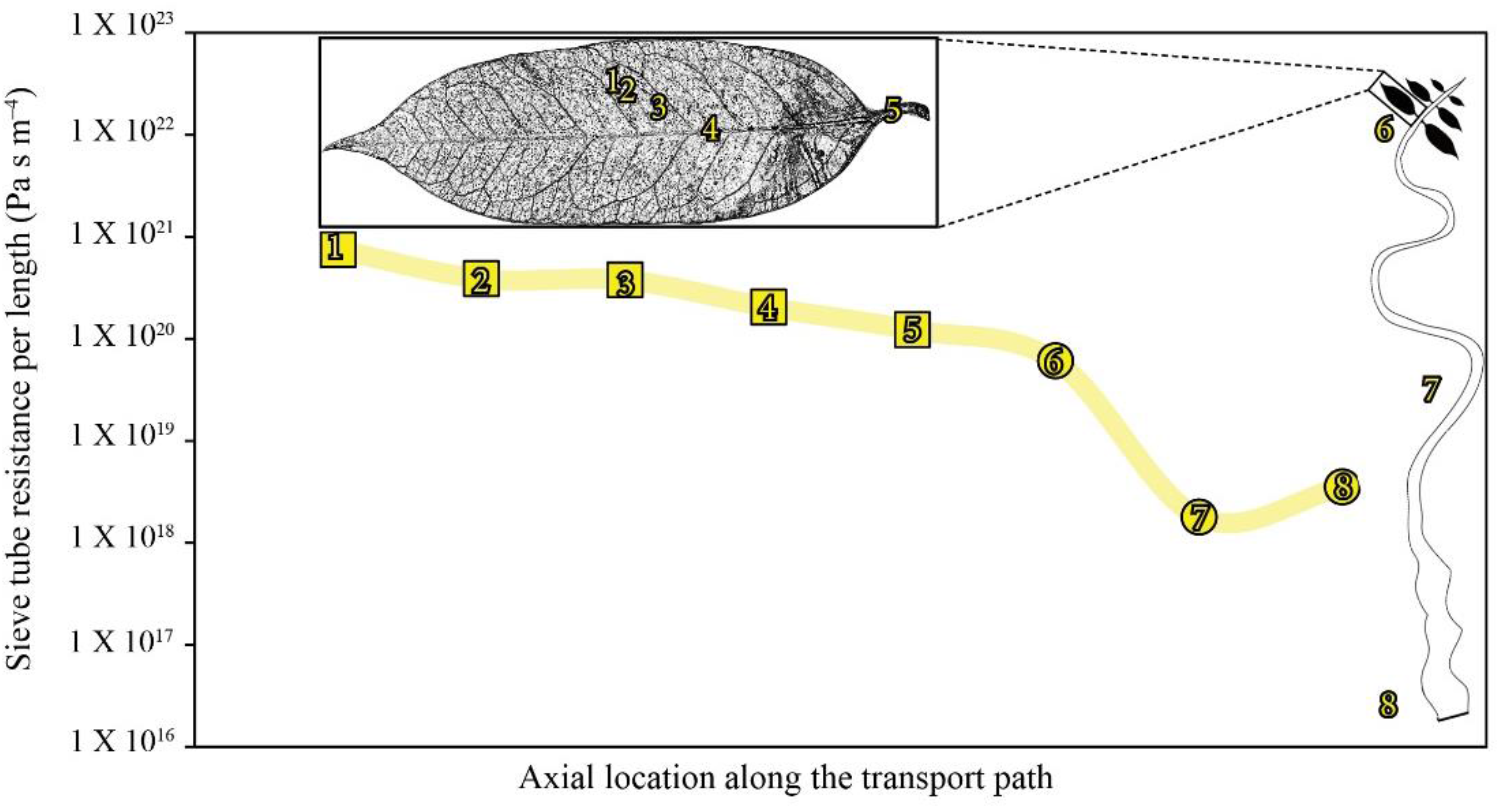
Sieve tube hydraulic resistance across the aerial organs of *A. scandens*. The sieve tube resistance drops two orders of magnitude from the minor leaf veins (squares) to the base of the stems (circles), even though the distance between them could be up to 20m. Numbers indicate the relative position along the transport pathway, noted in the figures, but they do not represent real distances.

## DISCUSSION

### PHLOEM ONTOGENY AND LIANESCENCE

During stem ontogeny, lianas that rely exclusively on twining for climbing, such as *A. scandens*, transition somehow abruptly from self-supporting branches, known as searchers, to stiffer ones that grow in close contact with a solid support. In *A. scandens*, the photosynthetic, non-lignified searcher branches displayed more phloem than xylem tissue, likely because the immature leaves have little demands of water, but a significant need of carbohydrates. This limited xylem implies that pith turgor might be the major contributor to the mechanical stability of searcher branches, as reported in other twining vines (Isnard and Silk, 2009). Morphologically, the tube-like primary sieve elements that dominated the vasculature of searchers suggested directional flow of nutrients for tip growth. Chemically, sieve tube walls were composed of a galacturonan-rich pectin, which has been described in a handful of species such as in leaves of *Beta vulgaris* (Torode et al., 2018), or in the seasonally renovated sieve tubes of *Populus* (Ray and Savage, 2020). These previous works strongly emphasized that the chemistry of the cell walls connect with mechanics of the sieve tube wall (Torode et al., 2018). Thus, the presence of this epitope likely contributes to the flexibility of searcher branches, but its concurrence in the primary sieve tubes of *A. scandens* further supports a conservation of this moiety in the sieve tubes of angiosperms with different life histories, growth habits, and phylogenetic backgrounds.

Rigid but slender stems displayed a lobed axial arrangement of the phloem tissue, as well as vermiform secondary sieve elements, with extremely angled sieve tube connections, both features suggestive of a morphology adapted to torsion. In fact, the complex division patterns of the secondary phloem of *A. scandens* were evaluated in detail about 50 years ago (Srivastava, 1970), and correlated with the helical gyres of the lianoid growth habit. We further observed that, in areas of high rotation, the sieve tube elements shortened, thus meeting the demands of helical growth (Silk and Holbrook, 2005). This reinforces the idea that the vascular cambium activity is influenced by the mechanics of lianoid stems (Pace et al., 2015, 2018). For example, it is widely accepted that the wider vessels evolved with the lianoid habit enhance the hydraulic efficiency, maintaining water supply to a dense canopy under strong torsional pressures such as twisting and bending (Putz and Holbrook, 1992; Gentry, 1991; Speck and Rowe, 1999; Rowe et al., 2006). Our measurements revealed that *A. scandens* has sieve tube geometries comparable to other woody species, such as the radii of sieve elements (7-13µm), which fall within parameters measured in the slender stems of *Ipomoea* (10-20µm) (Knoblauch et al., 2016), or the trunks of canopy dominant trees (6-24 µm) (Liesche et al., 2017; Savage et al., 2017). These diameters further follow escalation from thinner to thicker branches, as it is the case of tall trees (Jensen et al., 2012, Savage et al., 2017). Strikingly though, the average sieve plate pore size (0.20µm radius) was smaller than the stems of other species, and varied little axially. Because pore size is the factor that affects conductivity the most, *A. scandens* has lower phloem conductivity in the stem, compared with other species (Liesche et al., 2017; Savage et al., 2017; Clerx et al., 2020; Barceló-Anguiano et al., 2021a), but within the parameters observed in other Austrobaileyales of the understory (Losada and Holbrook, 2019). The secondary phloem of *A. scandens* evolved unusually high number of sieve plates along the tangential walls of phloem conduits. Compound sieve plates have been correlated with growth in height across woody species that reach the forest crowns (Pace et al., 2015; Knoblauch et al., 2016; Liesche et al., 2017; Savage et al., 2017; Losada and Holbrook, 2019; Clerx et al., 2020; Barceló-Anguiano et al., 2021a,b). The question remains on the mismatch between a highly conductive xylem and a phloem constricted hydraulically. Reasons behind may relay in the life history traits, such as the imbalance of carbon allocation between the leaf canopy and the stem tissues. Thus, volumetric xylem vessels meet hydraulic demands of wide leaf areas without the selective pressures of stem mechanical stability (Putz and Holbrook, 1992), but a reduced number of sink tissues does not require a more conductive phloem. Instead, mechanics of the twinning constrains the morpho-functional aspects of phloem transport in the understory organs.

### VASCULAR SCALING IN LEAVES OF AUSTROBAILEYALES AND OTHER ANGIOSPERMS

Within leaves, the geometry of vascular elements (tracheids and sieve tube elements) varied their dimensions across vein orders. Hierarchical scaling of the xylem has been extensively documented in the reticulate leaves of angiosperms, suggesting that the efficiency of water distribution across the continuous xylem conduits follows laws of energy conservation, such as Murray’s law (Murray, 1926; McCulloh et al., 2003; Carvalho et al., 2017a, Scoffoni et al., 2017). In contrast, the scaling of sieve tube elements in leaf veins of woody organisms remains poorly characterized. Recent works in leaves with different branching patterns, such as the dichotomously branched veins of *Ginkgo* (Carvalho et al., 2017b), or reticulated leaves of *Populus* and *Illicium* (Carvalho et al., 2017a; Losada and Holbrook, 2019), strongly suggest universal variation of the geometry of phloem conduits across leaf vein orders. Yet, an unexplored feature in the leaves of angiosperms is the size of pores connecting sieve tubes in the leaves. Our work provides novel evidence that pore sizes vary in accordance with sieve tube dimensions, from 0.08µm in minor veins to 0.12µm in the petiole. As a result, variation of both the geometry of tubes and size of pores contribute to building up a higher hydraulic pressure in the minor veins (loading tubes), which, along with viscous dissipation, enhance bulk export of photoassimilates toward the petiole under the pressure-flow predictions (Münch, 1930).

We further report a 1: 1.25 xylem to phloem ratio of areas of the major vein in *Austrobaileya* leaves, much more balanced in favor of the phloem than the general range of 1:4 to 1:10 reported in the leaves of deciduous trees (Artschwager, 1926; Waisel et al., 1966). More surprisingly, the isodiametric sieve tube elements in the midrib linearly increased in number toward the petiole. The dimensions of sieve elements in the major leaf veins remains poorly explored in angiosperms, but they are well documented in the single veined needles of conifers (Ronellenfitsch et al., 2015), in which photo assimilate export is favored by the increasing number of isodiametric phloem conduits from the tip to the base of the needles. This suggests a convergent strategy between the needles and the midrib, but whether this is conserved in other angiosperm leaves, needs further testing. Furthermore, the isodiametric tracheids of the midrib linearly increased in number toward the petiole, supporting predictions on the uniformity of xylem conduit diameter within the same vein order (McCulloh et al., 2003, 2009; Gleason et al., 2018). Among the scarce studies documenting xylem allometry in single veins, variation in conduit diameter along the midrib has been studied in *Fraxinus* (Petit et al., 2016) and *Acer* (Lechthaler et al., 2019). The shortening of tracheids toward the petiole of *A. scandens* revealed a constrained conductivity of water uptake by leaves, in sharp contrast with the highly conductive stems, pointing to hydraulic segmentation of the xylem between the stems and the leaves. In contrast, the sieve tube elements of the petiole reduced their length, but not their diameter, thus keeping a stable conductivity. We previously showed a similar trait in the leaves of *Illicium parviflorum* (Losada and Holbrook, 2019), but to confirm a possible conserved pattern across woody species, we compared these data with that of leaves from five temperate trees (*Acer saccharum, Liriodendron tulipifera, Catalpa speciosa, Liquidambar styraciflua*, and *Quercus rubra*). These evaluations revealed that, from the midrib to the petiole, sieve tube element radii were similar in all five species (Losada and Holbrook, unpublished data).

The special characteristics of the petioles have been previously put forward in a number of species such as in the genera *Beta* (Geiger et al, 1969), *Cyclamen* (Grimm et al., 1997), or *Pelagornium* (Ray and Jones, 2018), suggesting architectural plasticity that is uncoupled from the leaf lamina. Strikingly, we found that the petiole of *A. scandens* is fiberless, in sharp contrast with the massive presence of perivascular fibers in the leaf lamina. Fiberless petioles were further observed in *Illicium parviflorum* (Losada and Holbrook, 2019), and suggest that these short petioles require flexibility, especially in understory plants such as the majority of members of the *Austrobaileyales*, which need to orientate their leaves toward sun flecks. In addition, the petioles of *A. scandens* may aid with twinning, as previously suggested (Feild et al., 2003b), in agreement with previous evidence of a pivotal role of petiole reorientation in generating the squeezing force that stabilize twining stems (Isnard et al., 2009).

### CONTRASTING VASCULAR STRATEGIES BETWEEN LEAVES AND STEMS OF LIANAS IN THE UNDERSTORY

Our measurements, although limited in number, suggest that the velocity of the phloem in *A. scandens* is one order of magnitude slower than *Ipomoea* and in trees (Jensen et al., 2012; Knoblauch et al., 2016), but similar to the rates observed in *Illicium parviflorum* (Losada and Holbrook, 2019). While the radii of the phloem conduits were similar in petioles and searcher branches, the hydraulic resistance from leaves to stems followed a continuum, differing by three orders of magnitude from the minor veins to the base of the stems. Yet, this is a smaller difference than that reported between the top and the bottom of tree stems of comparable heights (Savage et al., 2017). Our calculations, which assume a constant viscosity of 1.7mPa s^-1^, would not allow transport at long distances with faster velocities. For example, the pressure required to transport the sap 3m from the leaves would be 1.4MPa, but longer distances would impair transport. In conclusion, with constant viscosity and velocity, the export of carbohydrates from the leaves is facilitated by the geometry of the phloem, but encounter architectural limitations in the stems (i.e. tiny pores). What this may imply is that the export of photoassimilates may easily be redirected toward the continuously growing canopy. This is possible because the stem girth is maintained for long distances, but enlarges typically at the base of the vine. Additionally, the stems of *A. scandens* are photosynthetic all along, pointing to the possibility of self-sustain, at least partially, the limited secondary growth, and root elongation.

The vesselless leaves of *A. scandens* were previously suggested as having a simpler anatomy than other angiosperms, such as the absence of palisade parenchyma, low stomatal density with slow responses to VPD changes (Feild et al., 2003), which correlated with a physiology associated to the understory environment (Brodribb and Feild, 2010). Similarly, woody members of the ancestral ANA grade, such as *Amborella trichopoda*, the only living member of the Amborellaceae, the family that shares an ancestor with all angiosperms, is vessel free and lacks reaction wood. (Feild et al., 2005; 2012). Altogether, these results that include a strong hydraulic segmentation of the xylem, and a high resistance of phloem to sap transport, correlate with the slow growth rate of this species in the understory conditions (Feild et al., 2012).

Exploring the phloem of extant members of the ancestral angiosperm grade ANA (Amborellales, Nymphaeales, Austrobaileyales) is particularly relevant, given that fossils, when available, rarely preserve this tissue. Thus, living members allow for the inference of the varied hydraulic solutions evolved -and perhaps maintained-by angiosperms during their initial radiation. *A. scandens*, the only extant member of the Austrobaileyaceae family, itself one of the three lineages composing the ANA grade (Amborellales, Nymphaeales, Austrobaileyales), a sister grade to all flowering plants (Mathews and Donoghue, 1999; Parkinson et al., 1999; Qiu et al., 2000; Soltis et al., 1999, 2018), has been historically used to speculate on the primitive growth habits of the first angiosperms. In particular, one of the critical questions is when and how flowering plants reached the canopy (Judd et al., 2018). Our work provides the first empirical interpretation of long distance transport in woody lianas with simple body plans, and the constraints associated with their life histories. Woody lianas, semi climbers and shrubs (rarely trees) dominate the extant forms of the earliest angiosperm lineages Amborellales and Austrobaileyales. Woodiness, considered the symplesiomorphic condition of angiosperms as a whole, was likely required to colonize the vertical niche during the Cretaceous, previously dominated through millions of years by gymnosperms. The question remains as to whether the great anatomical diversity of climbers, typically considered as highly derived, could have been the ancestral condition of angiosperms.

## ACKNOWLEDGEMENTS

We are grateful to William E. (Ned) Friedman for kindly sharing plant material and lab facilities and to the staff of the Arnold Arboretum of Harvard University for continuous support. This work was funded by the National Science Foundation IOS 1456845 research grant to N.M.H. Z.H. was awarded with a research internship program from the DaRin Butz Foundation at the Arnold Arboretum of Harvard University. JML is a ComFuturo researcher at the IHSM La Mayora, funded by FGCSIC, a RTI2018-102222-A-I00 grant from the Spanish Ministry of Science and Universities, and a LINKB20067 from CSIC.

## SUPPLEMENTARY MATERIAL

**Supplementary material 1.**
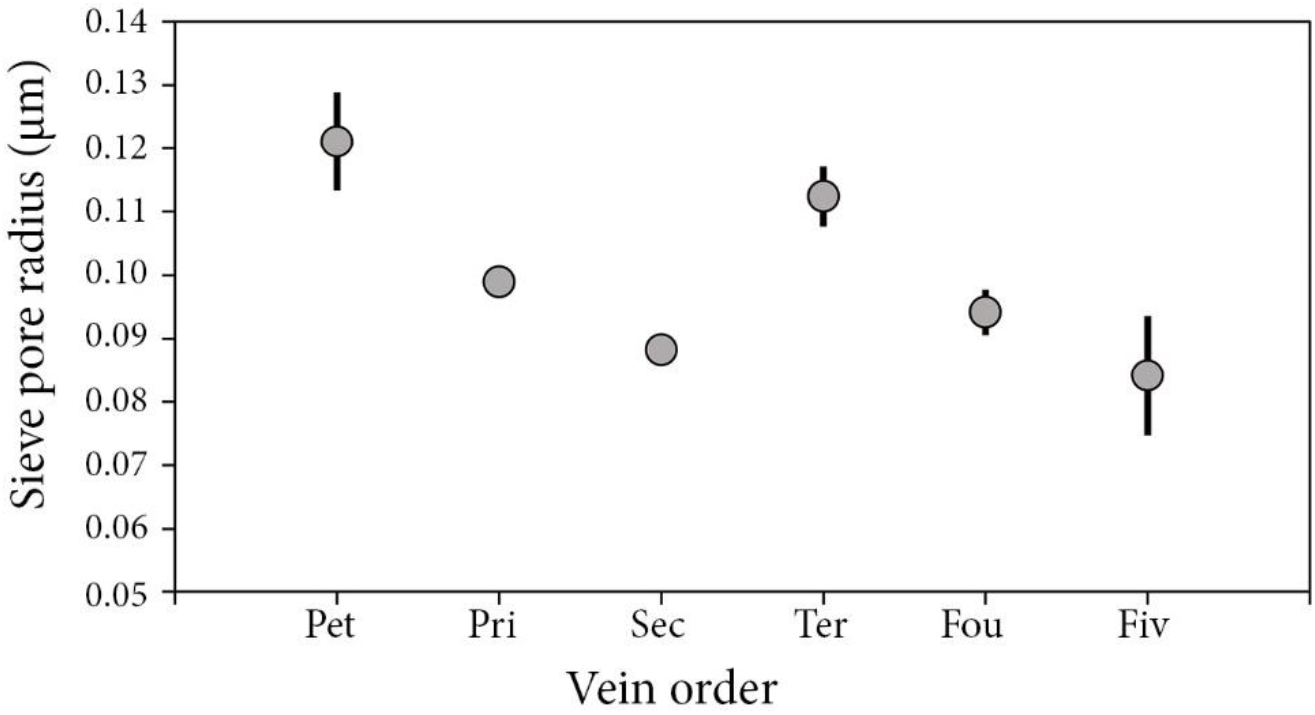
Pore radii in the leaves of A. scandens. X axis display the different vein orders: pet, petiole; pri, primary vein; sec, secondary vein; ter, terciary vein; fou, fourth vein order; fiv, fifth vein order. Bars represent standard error at a p<0.05.

